# Reverse gingival venipuncture: a refined technique for serial blood collection in small rodents

**DOI:** 10.64898/2026.04.24.720739

**Authors:** Jaturun Luapan, Haley Johnson, Melissa Seiberlich, Cory Brayton, Isabel A. Jimenez

## Abstract

**Objective:** To assess the safety, efficacy, and repeatability of a novel blood collection technique, percutaneous reverse gingival venipuncture (RGV), across multiple rodent species, and to characterize the associated anatomy through dissection and histopathology.

**Methods:** Success rate and complications of RGV were evaluated at a private practice between December 2024 and September 2025 in client-owned chinchillas (*Chinchilla lanigera*) (n=102), guinea pigs (*Cavia porcellus*) (n=78), Syrian hamsters (*Mesocricetus auratus*) (n=32), dwarf hamsters (*Phodopus campbelli* and *P. sungorus*) (n=4), squirrels (*Callosciurus erythraeus, C. finlaysonii*) (n=7), prairie dogs (*Cynomys ludovicianus*) (n=2), a capybara (*Hydrochoerus hydrochaeris*) (n=1) and a Patagonian mara (*Dolichotis patagonum*) (n=1). An experimental study was conducted in laboratory rats (*Rattus norvegicus*) (n=5), Chinese hamsters (*Cricetulus griseus*) (n=12), and chinchillas (n=11) from February 2026 to March 2026 to evaluate RGV success rate, serial hematology, and histopathology.

**Results:** The success rate of RGV was 100% in most rodent species, but was lower in guinea pigs (44.87%, n=78) and chinchillas (64.60%, n=113). No animals experienced clinical complications. No significant changes in serial hematology were appreciated in Chinese hamsters (n=6) on days 0, 7, and 14. Histopathology did not reveal any complications.

**Conclusions:** RGV in anesthetized rodents is safe, effective, minimally invasive, and repeatable, yielding clinically relevant blood volumes with precise control and minimal risks.

**Clinical Relevance:** RGV may facilitate more routine hematology and chemistry analysis in rodents by veterinary practitioners, with few risks and complications. In research settings, RGV may improve animal welfare and contribute to refinement and reduction.

## Introduction

Venipuncture is an essential technique for collecting blood samples for diagnostic and research purposes. Small rodents such as mice, rats, hamsters, chinchillas, and guinea pigs have grown significantly in popularity as pets in recent years,^1,2^ requiring more veterinarians to become comfortable managing these patients, including the ability to reliably collect diagnostic blood samples. Mice and rats are also the most common species used in research settings worldwide, with blood collection being a very frequent procedure performed by researchers for data collection. However, the small body size of most rodents poses an inherent challenge to reliable blood collection as it makes venous access technically challenging, limits sample volumes, and increases the risk of complications such as hemorrhage, tissue trauma, or mortality.

The route and methodology of blood collection has been shown to significantly affect stress levels, behavior, and tissue injury in rodents.^3–5^ Selection of an appropriate venipuncture route requires consideration of desired volume yield, intended purpose of the sample, frequency of serial collection, and requirement of anesthesia. A strong understanding of species-specific anatomy, physiology, and behavior is also essential to optimize the chance of success. Further, the sample volume that can be safely collected is dictated by the patient’s circulating blood volume. The general guidance is not to collect more than 10-15% of circulating blood volume at one time to preserve hemodynamic stability, and to allow 2-4 weeks between collection of this maximum volume to allow time for the replenishment of blood components.^6,7^ Smaller volumes can be collected more frequently. However, in small rodents, sample volumes required for diagnostic or research assays may approach the upper limit of what can safely be collected at one time. Certain collection routes can lead to hemorrhage, such as the “cheek puncture” in mice which involves puncturing the facial vessels, and must be factored into the total collection volume.^8,9^ Routes that allow more precise control of sample volume are thus advantageous, and in research settings may enable serial blood collection and facilitate non-terminal longitudinal studies, offering a refinement and reducing animal numbers. Blood collection methods for small rodents that reliably yield adequate sample volumes and limit hemorrhage and tissue trauma, while being simple to train and perform, are thus necessary in both clinical and research settings.

Common sites for venipuncture in rodents include the lateral tail vein, facial vein, saphenous vein, external jugular vein, cranial vena cava, lingual vein, and retro-orbital sinus/plexus. Advantages and risks for these routes in each species have been summarized in **Supplementary Table S1**. References are included where available, and for some routes, additional risks are also included by the authors of the present study based on experience in both clinical and research settings. It should be noted that for several of these techniques, multiple names have been associated with the target vessels and/or anatomical approaches, which can lead to confusion.^10,11^ For example, two studies referencing blood collection from paired vessels running along the ventral tongue describe these as sublingual veins.^12,13^ However, anatomical dissections suggest that the two vessels are more appropriately termed the lingual veins (*v. lingualis*).^14,15^

In general, peripheral vessels such as the lateral saphenous and cephalic veins, which are commonly used venipuncture sites in other mammals, are small and fragile in rodents and yield small sample volumes. Blood collection from peripheral vessels can be facilitated by preheating the are with a heat lamp, warm water, or warming pad, or by using a tourniquet. However, care must be taken to avoid overtightening or prolonged placement of the tourniquet, as this could lead to tissue injury. Several routes that yield larger blood volumes, such as the retro-orbital sinus in mice, retro-orbital plexus in rats, or the cranial vena cava in guinea pigs and chinchillas, carry higher risks of complications and require anesthesia to ensure immobility and prevent pain or distress. Other approaches that yield large volumes may only be performed as non-survival procedures, such as terminal cardiocentesis, rendering these techniques applicable in some research contexts but not useful in a clinical setting.

The gingival vein (*v. labialis mandibularis*), located along the rostral midline on the labial aspect of the root of the mandibular incisors, has more recently been described as a reliable site for venipuncture in small rodents, including rats, mice, guinea pigs and hamsters.^16,17^ This site is minimally traumatic and allows for repeated sampling with precise control of volume collected and relatively high yields.^17^ Notably, Rodrigues et al. (2017) found no significant changes in complete blood count (CBC) findings following serial collection from this site in guinea pigs and hamsters, and no histopathological evidence of trauma or inflammation to the gingival tissues. The gingival venipuncture technique described in the literature involves approaching through the mouth, from rostral to caudal, inserting the needle through the gingiva over the lower incisors at a 20-25º angle and encountering the vessel along midline, approximately 2 mm deep.^16^ In this approach, the needle passes through the gingiva before it enters the vessel, precluding preemptive disinfection of the site. In guinea pigs, bacterial contamination was common in samples taken via this route, although this did not lead to significant changes in serial CBCs.^18^ In addition, the traditional approach to the gingival vein is expected to result in the presence of blood within the oral cavity, which could be distressing to the patient upon recovery from anesthesia.

Here, we evaluate the safety, efficacy, repeatability, and associated anatomy of a novel blood collection approach, termed percutaneous reverse gingival venipuncture (RGV). The designation “reverse” refers to the approach being opposite to that of the conventional gingival vein technique. The newly developed method approaches from caudal to rostral, where the interdental diameter is wider, with the needle passing through the skin followed by the mucosa before entering the target vessel. We hypothesized that in Chinese hamsters, three serial blood collection events via RGV of approximately 15% of circulating blood volume, at 7-day intervals, would not result in significant changes to hematology results over time. We predicted that the RGV technique would be simple to train for new operators, demonstrating the feasibility of implementing RGV in both clinical and research settings. We further hypothesized that the RGV technique would have a high rate of success, defined as a sample yield of ≥0.1 mL, with minimal or no complications in various rodent species, thus representing a refinement for blood collection in rodents.

## Materials and Methods

### Animals

A subset of animals in this study were client-owned animals undergoing anesthetized clinical veterinary evaluation and clinically indicated blood collection at Animal Space Pet Hospital, Bangkok, Thailand. These included 102 chinchillas (*Chinchilla lanigera*), 78 guinea pigs (*Cavia porcellus*), 32 Syrian hamsters (*Mesocricetus auratus*), 4 dwarf hamsters (*Phodopus* spp.), 7 squirrels (*Callosciurus erythraeus, C. finlaysonii*), 2 prairie dogs (*Cynomys ludovicianus*), 1 capybara (*Hydrochoerus hydrochaeris*) and 1 Patagonian mara (*Dolichotis patagonum*). Anesthesia was performed in most cases, as determined to be appropriate for each individual patient by the veterinarian (JL). In some cases, collection was able to be performed in conscious animals with brief manual restraint, particularly chinchillas. RGV was performed as described below, with needle gauge varying between 25-30G depending on the species.

A separate experimental study was conducted in rats (n=5), Chinese hamsters (n=12), and chinchillas (n=11) housed at Johns Hopkins University, Baltimore, MD. All work was conducted with ethical approval by the Johns Hopkins University Animal Care and Use Committee. Animals were maintained in accordance with federal regulations and the *Guide for the Care and Use of Laboratory Animals* in an AAALAC-accredited facility. Chinchillas were housed on an unrelated research protocol and underwent anesthesia for routine examinations and collection of blood for annual hematology and chemistry; no chinchillas were euthanized for the present study. Rats and Chinese hamsters used in this study were experimentally naïve (e.g. retired breeders) originally housed on unrelated research protocols, which were no longer required by the primary investigator and had already been designated for euthanasia. Thus, no new animals were acquired or purchased specifically for this study.

A total of 11 chinchillas underwent anesthesia with isoflurane inhalation (3% for induction, 2% for maintenance using a nose cone). RGV was attempted in all chinchillas as described below using a 25G or 29G needle. RGV was attempted by two operators for each chinchilla; operators were classified as “first-time operator” if they had never attempted RGV before, “new operator” if they had attempted RGV before but had not successfully performed two RGVs, and “trained operator” after they had performed two successful RGVs in this species. If RGV was unsuccessful after two attempts, blood was collected via the external jugular vein using a 25G needle.

Five laboratory rats were used in this study. Three rats underwent anesthesia with 40-80 mg/kg ketamine + 2.5-5 mg/kg xylazine IP, followed by 2-4 serial RGV attempts using a 29G needle attached to a 1 mL, U-100 insulin syringe (Sol-Millenium Medical Inc., Suwanee, GA 30024). One rat was euthanized by CO_2_ inhalation immediately after RGV for necropsy and histopathology. Two rats were administered 0.2 mg/kg atipamezole IM and recovered uneventfully from anesthesia, after which they were monitored for 7 days prior to euthanasia. Two control rats that did not undergo RGV were euthanized by CO_2_ inhalation and included for anatomical dissection and/or histopathology. Ten Chinese hamsters (*Cricetulus griseus*) were sedated with 40 mg/kg ketamine IP, which resulted in marked muscle relaxation but did not fully eliminate motor activity or eliminate righting reflex. Brief manual restraint with a firm scruff hold was used to properly position the hamsters for blood collection. RGV was performed using a 29G needle attached to a 1 mL, U-100 insulin syringe, and 15% of circulating blood volume, estimated as 10% of the animal’s body weight (0.25-0.4 mL depending on body weight) was collected for CBC. Immediately after RGV, 4 hamsters (2 male, 2 female) were euthanized for necropsy and histopathology, while the remaining 6 hamsters underwent additional RGV events at day 7 and day 14 for serial hematology analysis and comparison to reference values.^19^ Two additional Chinese hamsters that did not undergo RGV were euthanized by CO_2_ inhalation for necropsy and histopathology.

### RGV technique

Animals were positioned in dorsal recumbency with the neck straightened. The skin was disinfected with alcohol. The triangular region at the base of the mandibular incisors, just below the mandibular symphysis, was palpated to identify the landmark for insertion. A 25-30G needle, selected according to species, attached to a 1 mL syringe was introduced along midline at this location, passing through the skin and into the mucosal region. The angle of placement varied between 0º to 60º depending on the species. Once the bevel of the needle was beneath the skin, gentle negative pressure was applied to the syringe while the needle was slowly advanced until blood entered the needle hub, after which gentle and consistent negative pressure was maintained until the sample was collected. The depth of the needle was closely monitored to ensure that the needle did not enter the oral cavity. Manual pressure was applied after the needle was withdrawn and maintained until hemostasis was confirmed. For this study, the success rate of RGV was defined as a sample yield of ≥0.1 mL.

### Hematology

Blood samples from chinchillas were deposited directly through the needle into commercial potassium EDTA blood tubes (BD Microtainer, BD, Franklin lakes, NJ 07417). Chinchilla hematology samples were run by a commercial diagnostic laboratory (IDEXX BioAnalytics, North Grafton, MA 01536). After collecting blood from Chinese hamsters, the plunger of the syringe was drawn back to create space and the barrel was cut to remove the needle, then the blood sample was deposited into potassium EDTA blood tubes. Chinese hamster hematology was analyzed in-house using the Procyte Dx (IDEXX Laboratories, Westbrook, Maine 04082). Chinese hamster hematology results were analyzed in GraphPad Prism 11.0.0, with changes in hematology parameters evaluated over time via one-way repeated measures ANOVA with Geisser-Greenhouse correction, and Dunnett’s post hoc test for multiple comparisons. Hematology parameters were also compared between male and female Chinese hamsters via a two-way repeated measures ANOVA with Geisser-Greenhouse correction.

### Histologic evaluation in rats and hamsters

After euthanasia of rats and hamsters, anatomical dissection of the head and neck region was performed. In four rats, mandibles with skin and tongue were collected intact, then fixed and decalcified in Formical4 (Statlab, McKinney, TX 75069) for 7 days. Mandibles were either trimmed to transverse sections from the labial to the glossal aspect of the incisors or divided midsagittally, then processed routinely, including paraffin-embedding, sectioning at 5 µm, and stainined with hematoxylin and eosin (H&E). Slides were scanned using the Zeiss Axio Scan.Z1 platform (Zeiss, Jena, Germany) and viewed on Concentriq LS software (Proscia, Philadelphia, PA 19103). Labeled figures were prepared using Adobe Photoshop 46.4.0.

## Results

The anatomic landmarks and angle of approach in different rodent species are shown in **Figures 1-3**. The combined success rates for RGV in both the clinical and experimental studies revealed that RGV attempts were successful in 100% of Chinese hamsters (n=22), dwarf hamsters (n=4), rats (n=4), prairie dogs (n=2), capybara (n=1), and Patagonian mara (n=1). Success rates were high in Syrian hamsters (93.75%, n=32) and squirrels (85.71%, n=7), but were lower in chinchillas (64.60%, n=113) and guinea pigs (44.87%, n=78). In a few cases, RGV was able to be performed under brief manual restraint, particularly in chinchillas; however, most patients required sedation or anesthesia.

**Figure 1.**
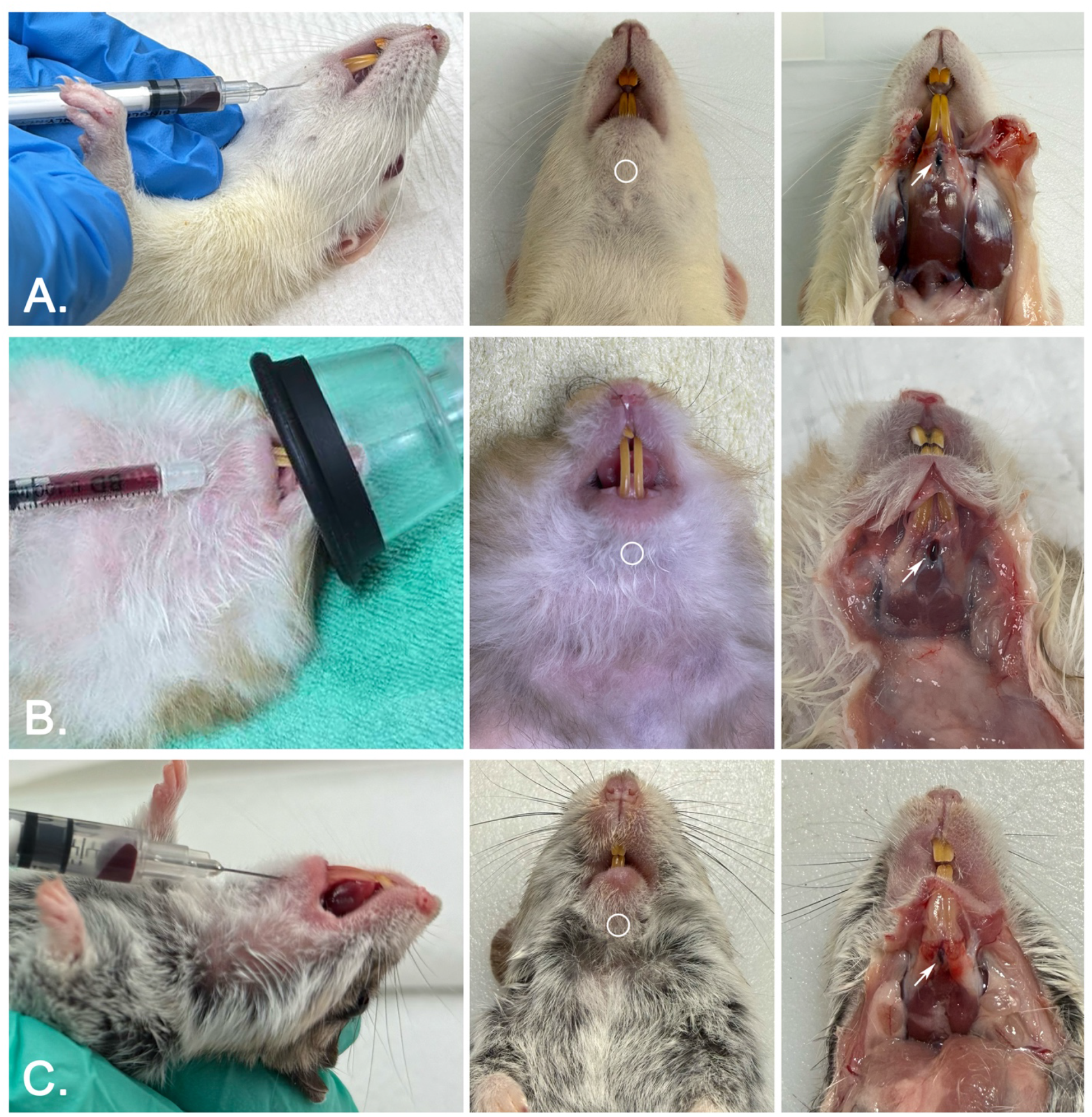
Anatomical characterization of the gingival vein and reverse gingival venipuncture (RGV) landmarks. Circle indicates target site of insertion of needle. White arrow indicates the gingival vein. **A)** Rat. **B)** Syrian hamster. **C)** Chinese hamster.

**Figure 2:**
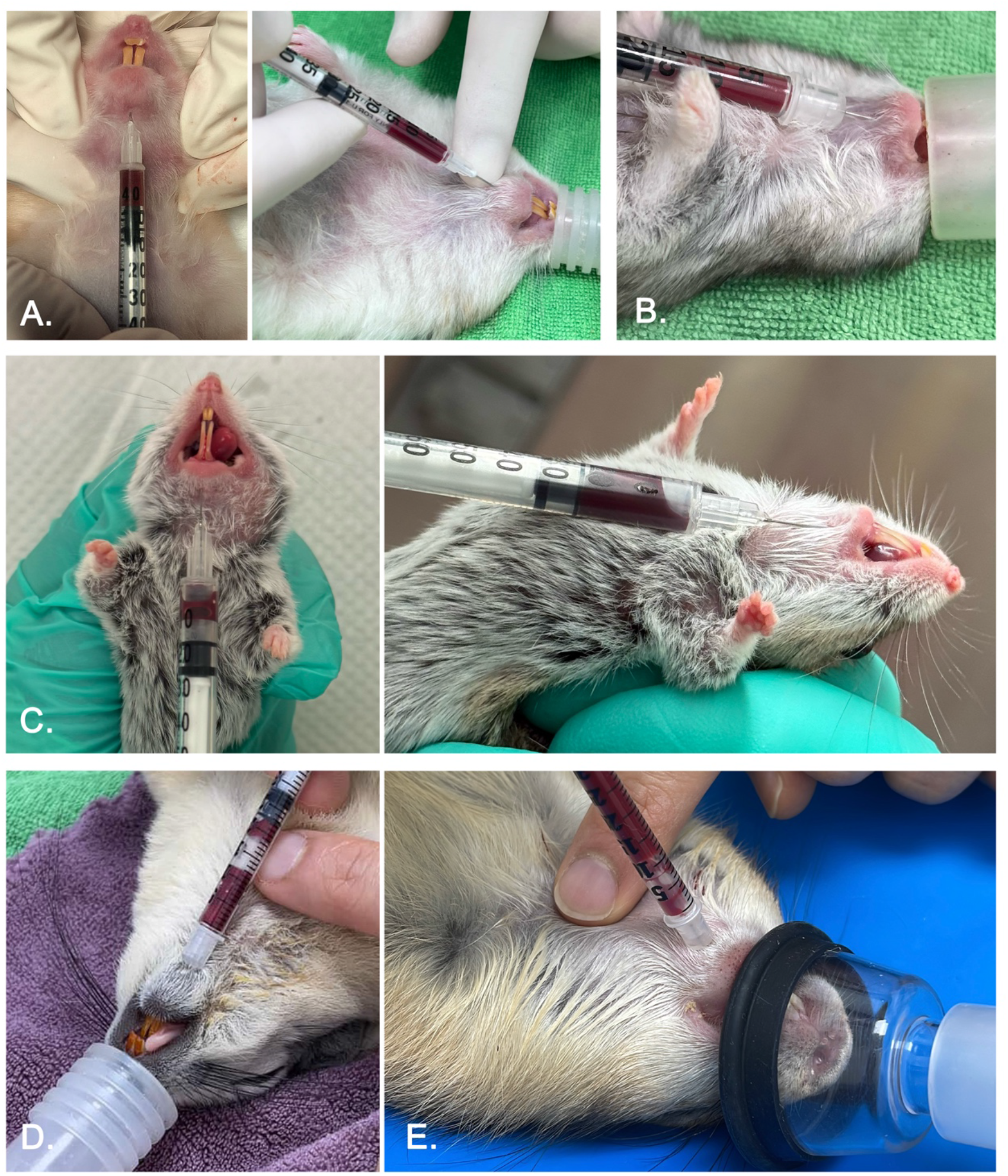
Reverse gingival venipuncture (RGV) technique in hamsters and Sciuromorpha. With the rodent in dorsal recumbency, the triangular landmark at the base of the mandibular incisors is palpated and a 29G needle is inserted. Once the bevel is below the skin, gentle negative pressure is applied while the needle is advanced until blood is present in the hub. The angle relative to the neck is adjusted depending on the species. The depth of the gingival vein varies between rodent species; for small rodents such as hamsters, the vessel can be encountered approximately 2 mm beneath the skin, while in larger rodents, the needle may need to be advanced further. To reduce the risk of hemolysis, slow gentle aspiration is utilized, and the needle is removed manually or by cutting the barrel of the syringe before placement of blood into the collection tube. **A)** Syrian hamsters (30-45º angle). **B)** Dwarf hamsters (20-30º angle). **C)** Chinese hamsters (0-30º angle). **D)** Squirrel (30-45º angle). **E)** Prairie dog (30-45º angle).

No hemorrhage or other adverse clinical effects occurred in association with RGV in this study and in client-owned animals, no owners reported any complications following discharge from the hospital. Histopathology in rats (n=1) and hamsters (n=2) showed perivascular hemorrhage immediately after blood collection, but no abnormalities 7 days later (n=2), including no evidence of necrosis, inflammation, trauma to the dental germinal tissue or alveolar bone, or infection associated with the soft tissues, alveolar bone, or teeth.

In the research assessment with 11 chinchillas, the success rate for a first-time operator performing RGV was 50% (3/6), while the success rate for a trained operator was 83% (5/6). The success rate for a new operator ranged from 0-100%. In Chinese hamsters, RGV was successful in trained operators in 100% of collection events (n=24) with sample yield ranging from 0.1-0.4 mL.

In Chinese hamsters, serial CBCs showed no significant change over time in hematocrit (HCT) (P=0.1592), hemoglobin (P=0.3784), platelets (PLT) (P=0.4943), total white blood cell (WBC) count (P=0.2030), heterophil count (P=0.3306), heterophil % (P=0.5306), lymphocyte count (P=0.1669), and lymphocyte % (P=0.5686). There were also no significant differences between male (n=3) and female (n=3) Chinese hamsters with regards to HCT (P=0.2787), hemoglobin (P=0.3238), PLT (P=0.1269), WBC (P=0.3346), heterophil count (P=0.1462), and lymphocyte count (P=0.5197). There was a significant effect of sex on heterophil % (P=0.0103) and lymphocyte % (P=0.0083), with females having a significantly higher percentage of heterophils and males having a significantly higher percentage of lymphocytes. There was also a significant effect of individual hamster on total WBC count (P=0.0081), neutrophil count (P=0.0085), and lymphocyte count (P=0.0173). Bloodwork results are shown in **Supplementary Figure S1**.

In our study, chinchilla blood samples collected with 25-29G needles were not removed from the syringe prior to depositing the blood into the collection tube. Of these, 3/3 samples collected by RGV and 3/4 samples collected from the external jugular vein had hematocrits within the normal reference range and did not show evidence of hemolysis, while 1 sample from the jugular vein was hemolyzed. Chinese hamster blood samples were collected with a 29G needle which was then removed by cutting the barrel of the syringe with canine nail clippers (Millers Forge, Plano, TX, 75074) prior to dispensing the blood into the collection tube, and no samples exhibited hemolysis.

On gross anatomical dissection in the rat, Syrian hamster, and Chinese hamster, a large vessel was visible superficially along midline between the roots of the mandibular incisors, just below the level of the mandibular symphysis (**Figure 1**). On gross dissection in the rat, once the mandibular symphysis was divided, two gingival veins were appreciated, adjacent to each other at the symphysis and then running along the medial aspect of the mandibles. Small-caliber paired lingual veins coursed parasagittally beneath the tongue. No appreciable vessels or vascular spaces were identified in the floor of the mouth in the rat. On histopathology of rats (**Figure 4**) and Chinese hamsters (**Figure 5**), gingival vein(s) were visible close to midline, <2 mm deep, and larger, thin-walled vasculature structures, compatible with sinuses, were identified medial to each mandible. In the Chinese hamster, the lingual vein appeared to drain into this sinus.

**Figure 3:**
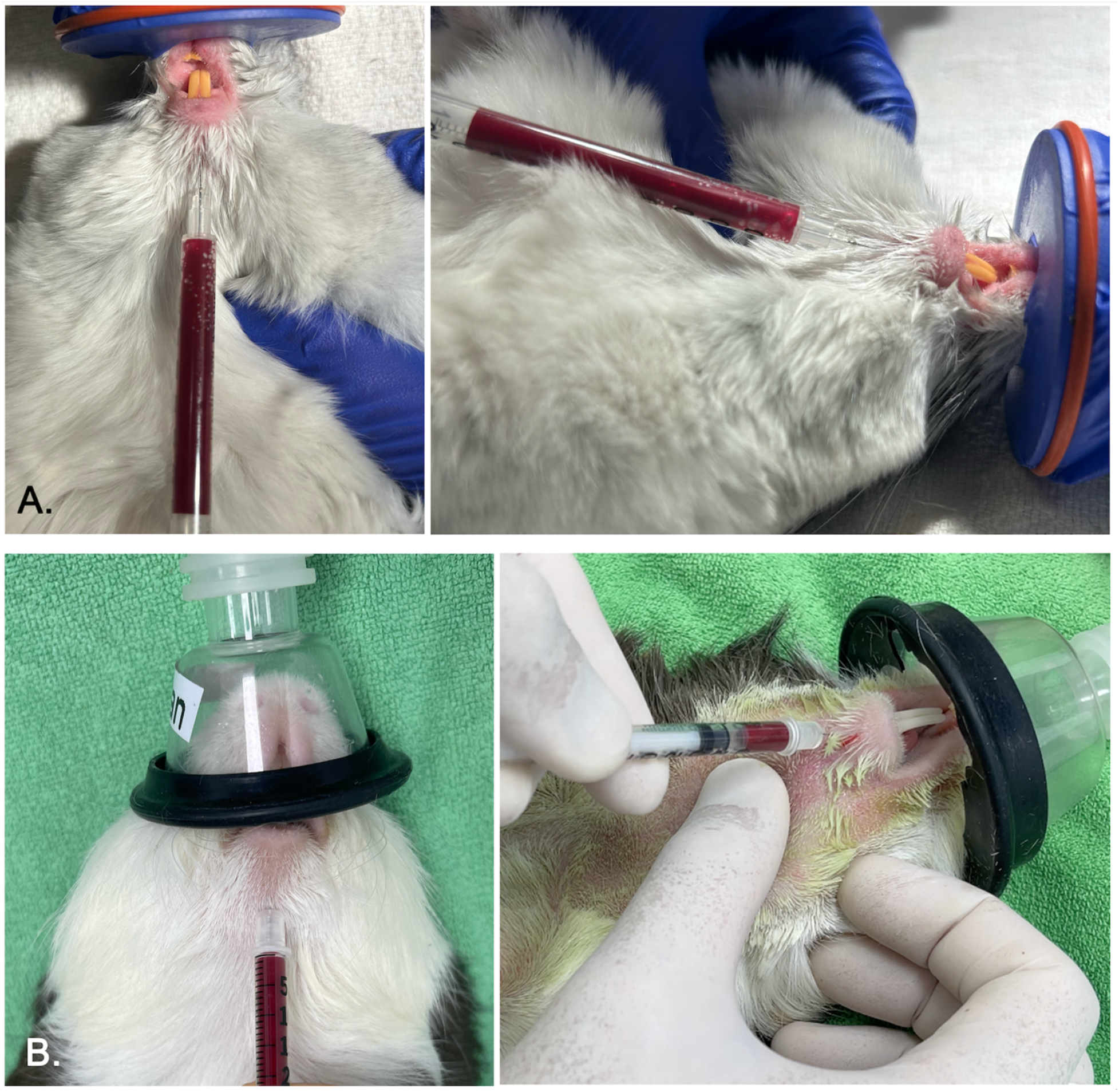
Reverse gingival venipuncture (RGV) technique in Caviomorpha. With the rodent in dorsal recumbency, the triangular landmark at the base of the mandibular incisors is palpated and a 29G needle is inserted. Once the bevel is below the skin, gentle negative pressure is applied while the needle is advanced until blood is present in the hub. The angle relative to the neck is adjusted depending on the species. The depth of the gingival vein varies between rodent species. In chinchillas, the vessel can be encountered approximately 2 mm beneath the skin, while in guinea pigs, the vessel was deeper. To reduce the risk of hemolysis, slow gentle aspiration is utilized, and the needle is removed manually or by cutting the barrel of the syringe before placement of blood into the collection tube. **A)** Chinchillas (20-30º angle) **B)** Guinea pigs (30-45º angle)

**Figure 4.**
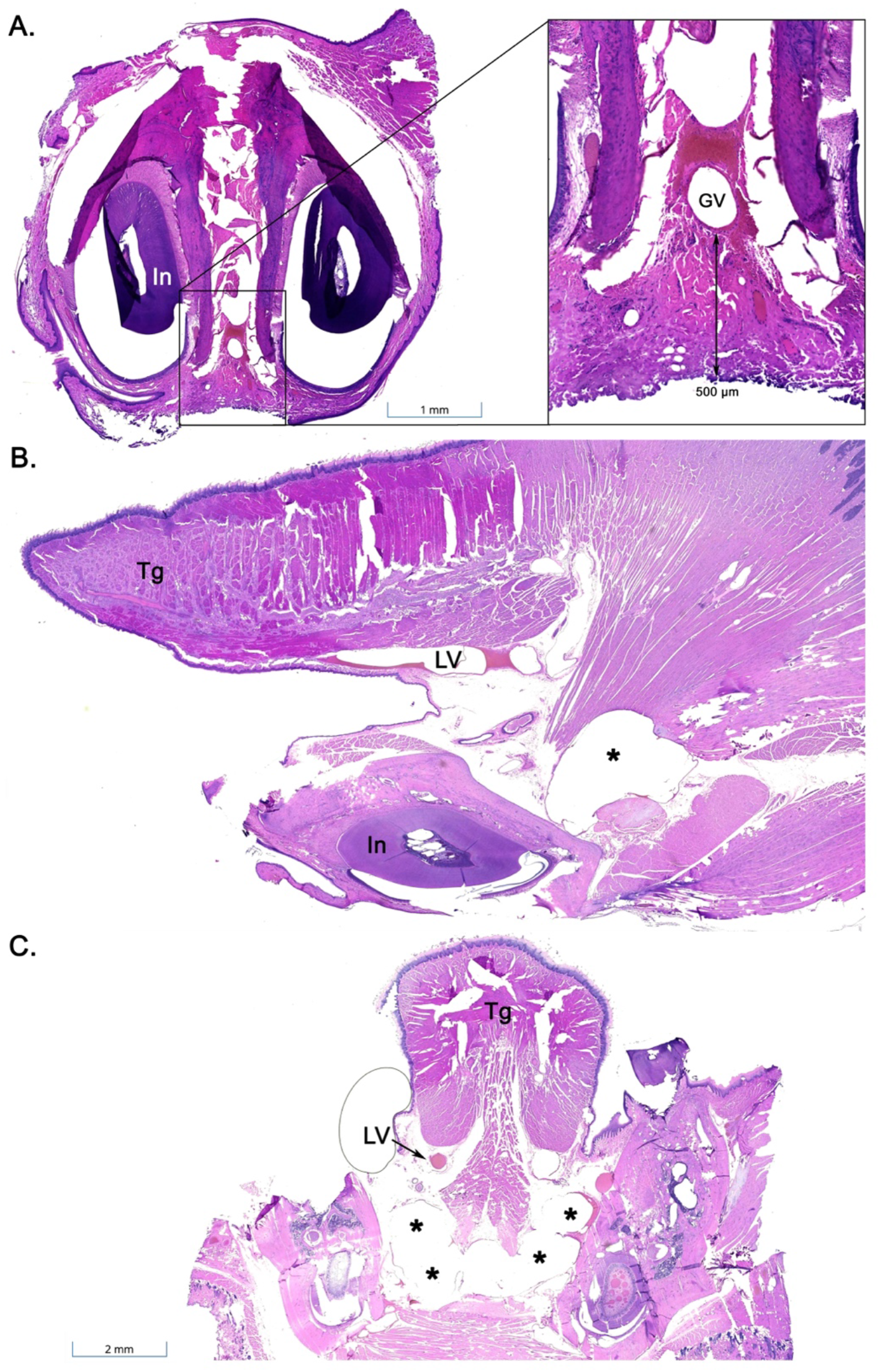
Histopathology of the gingival vein and identification of a large vascular space between the mandibles of the rat. A) Transverse section of the rat mandible, with a gingival vein (GV) located submucosally between the mandibular incisors (In). B) Parasagittal section of the rat mandible, showing the lingual vein (LV) ventral just to the tongue (Tg), and a thin-walled vascular space (*) caudal to the incisors (In). C). Transverse section of the rat mandible taken more caudally than B., with the lingual veins (LV) visible ventral to the tongue (Tg) and large bilateral vascular spaces (*). H&E.

**Figure 5.**
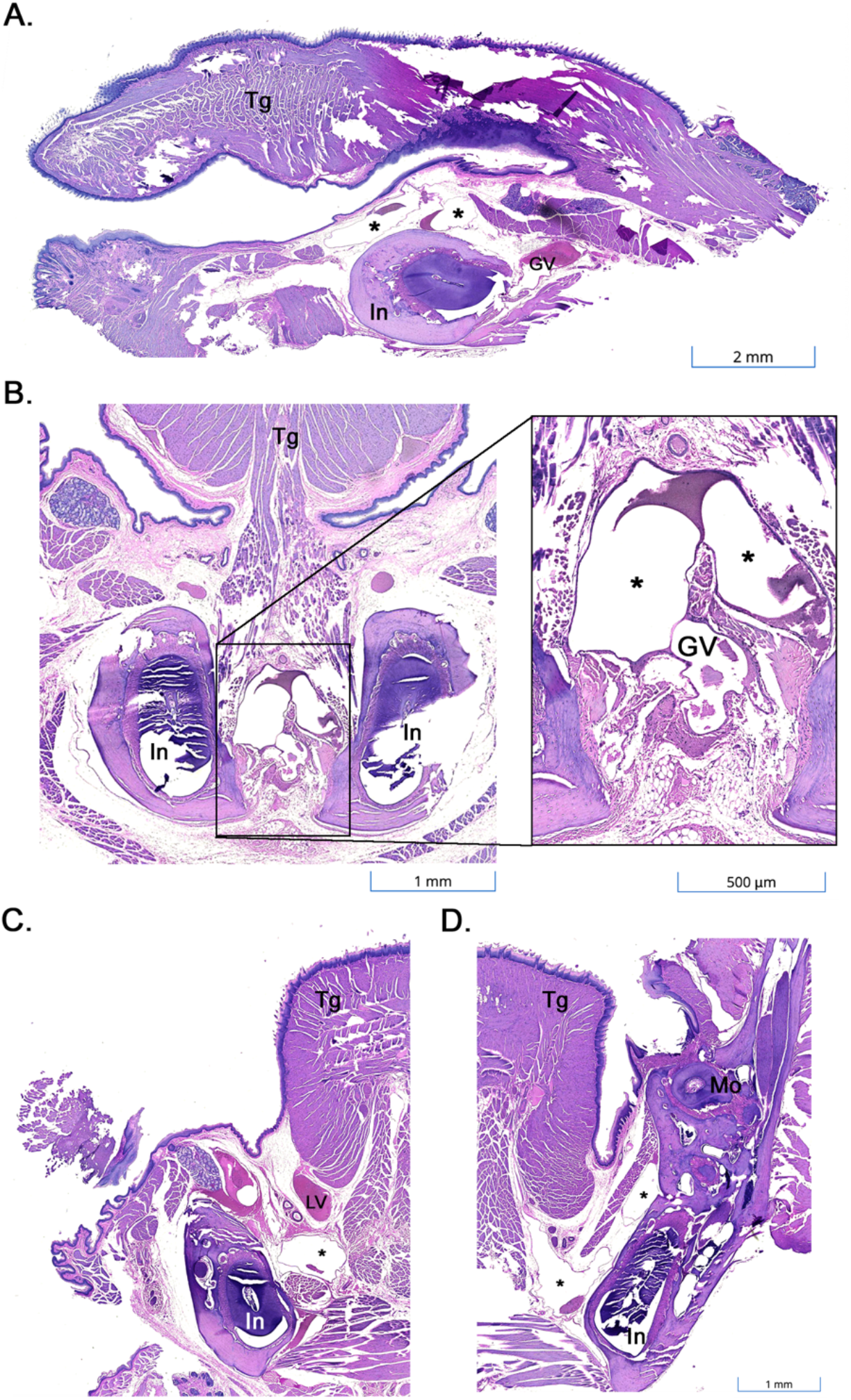
Histopathology of the gingival vein and identification of a large vascular space between the mandibles in the Chinese hamster. A) Parasagittal section of the Chinese hamster mandible, with a gingival vein (GV) located submucosally between the mandibular incisors (In) and a large, thin-walled vascular space (*). B) Transverse section of the Chinese hamster mandible, with a gingival vein (GV) and bilateral vascular spaces (*) located between the incisors (In). C) Transverse section taken more caudally than B., with the lingual vein (LV) ventral to the tongue (Tg) and a large thin-walled vascular space medial to the incisor (In). D) Transverse section taken more caudally than C., with large, thin-walled vascular structures medial to the mandible. A vein within the tongue appears to drain into this sinus. The roots of the incisor (In) and a molar (Mo) are also shown. H&E.

## Discussion

The RGV approach was a feasible, safe, and effective blood collection technique across multiple rodent species, allowing precise volume control and minimal risk of complications. RGV was highly successful in most species evaluated, but was more challenging in chinchillas and guinea pigs, which may be due to their narrower interdental region and copious loose skin associated with the chin and lower lip. While there was variability success rate for first-time or new operators, after a brief period of training, proficiency was gained rapidly and the success rate for trained operators performing RGV in Chinese hamsters and chinchillas was 100% and 83%, respectively. All personnel performing venipuncture in the current study were licensed veterinarians. Further studies investigating the teachability of this technique for researchers without specialized technical training on venipuncture via other routes are necessary to fully evaluate its potential for implementation in research settings. H

While RGV could be performed under brief manual restraint in some patients, particularly in chinchillas, conscious RGV should be performed only by proficient operators and with careful consideration of species and individual patient factors. The authors recommend that anesthesia be used in most cases. For example, the stocky bodies of guinea pigs and the copious loose skin of hamsters make it challenging to restrain conscious animals in a way that entirely prevents jaw movement. Since the gingival vein is very superficial, care must be taken not to over-advance the needle during RGV, which in smaller animals, could lead to trauma to the oral cavity. We anticipate that this risk would be increased in a conscious patient that could move their jaw during blood collection.

In Chinese hamsters, serial hematology showed no significant changes over time, consistent with the findings of Rodriguez et al. (2017) using the conventional gingival venipuncture technique. Further studies, with larger sample sizes, that assess hematologic parameters in other rodent species over time and compare these to other documented blood collection techniques are also recommended to evaluate the value of RGV as a refinement.

Based on the morphological structures of the mentolabial region, variations in the interdental space and depth of the gingival vein among rodent species may lead to differences in technique and blood collection quality. Determination of species-specific optimal sites requires further investigation in future studies. In our study, collecting serial samples via RGV was consistently feasible in all species. From the authors’ experience, rodents of the Myomorpha group (“mouse-like” or “rat-like”), including rats and hamsters, appeared to exhibit higher venous pressure during RGV compared with those of the Sciuromorpha (“squirrel-like” including squirrels, prairie dogs) and Caviomorpha (guinea pigs, chinchillas), as assessed by the ease with which blood flowed into the syringe with negative pressure. In addition, Caviomorpha have a relatively narrow interdental region for their body size, therefore smaller needles are recommended.

A critical consideration for RGV is the potential for hemolysis associated with small needles and high negative pressure. To mitigate this risk, gentle but consistent aspiration is essential. In addition, we recommend removal of the needle following collection to preventing hemolysis associated with deposition of the sample into the collection tube. In the case of tuberculin or insulin syringes where the needle cannot be manually removed, the barrel of the syringe can be cut. If hemolysis is present, BUN, creatinine, amylase, and lipase may still be interpreted, while sodium, potassium, phosphorus, and total protein are rendered unreliable in hemolyzed samples.^20^

To the authors’ knowledge, the thin-walled vascular spaces that we visualized on histopathology in rats and Chinese hamsters have not previously been identified in rats or hamsters. The anatomy of the vasculature of the lower jaw, incisors and gingiva in rodents remains to be fully characterized. Textbooks on the anatomy of small rodents do not remark upon the gingival vein, nor do they identify a nearby venous sinus, plexus, or vascular space.^15,21^ A submental plexus has been described in rats, located bilaterally at the lateral aspect of the mandible near the alveolus of the incisor and draining into the inferior labial vein (*v. labialis inferior*).^15^ The inferior labial vein then converges with the facial vein (*v. facialis*) on each side.^10,15^ Given the lateral location, however, this submental plexus does not appear to correlate with the thin-walled vascular structures we identified on histopathology in the present study. A submental plexus has also not been described in mice, hamsters, or guinea pigs.^15^ Several other studies focus on the large vessels of the head and neck, but do not assess the vasculature at the base of the incisors.^22–24^ Future studies should prioritize anatomical dissection to further characterize the gross anatomy of the thin-walled structures here identified on histopathology.

The focus of this study was to test the feasibility, repeatability, and adverse effects of a novel blood collection technique in a variety of small rodent species, and to identify the relevant associated anatomy. We demonstrate that RGV is an attractive venipuncture technique in small rodents that is minimally invasive, safe with minimal risk of hemorrhage, and an effective collection method for serial sampling of large blood volumes, with a high success rate across many species. RGV may be particularly useful for guinea pigs, chinchillas, and hamsters, given that peripheral vessels are challenging to access due to their stocky bodies, short limbs, and short tails. This novel venipuncture technique also has benefits over the traditional gingival method because it allows sanitation of the skin prior to collection, eliminates oral mucosa contamination, avoids leaving blood within the oral cavity, and can be performed by a single operator. RGV thus represents a refinement to blood collection in small rodents.

## Acknowledgements

The authors thank Maria Maggio, VMD and Kelly E. Lane, LVT, for their assistance with anesthesia in rats and chinchillas.

