## Supplementary Figure S1 for "Reverse gingival venipuncture: a refined technique for serial blood collection in small rodents"

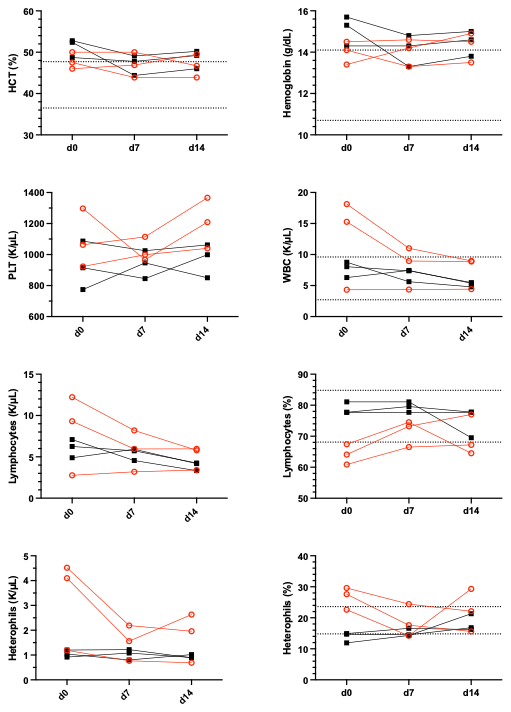

**Supplemental Figure 1:** Bloodwork values from serial CBC. Female hamsters are indicated by red circles and male hamsters are indicated by black squares. Reference ranges are indicated with dotted lines.^50^
