## Supplementary Table S1 for "Reverse gingival venipuncture: a refined technique for serial blood collection in small rodents"

**Supplementary Table S1: Common routes of non-terminal blood collection in small rodents**

| **Species** | **Site** | **Anesthesia required*** | **Volume** | **Potential Risks** | **References** |
| --- | --- | --- | --- | --- | --- |
| **Mouse** | “Cheek puncture” aka submandibular route (targets facial, superficial temporal, maxillary, or linguofacial veins) |  | 0.1-0.2 mL | Hematoma  Hemorrhage  Facial trauma  Trauma to ear canal  Bacterial contamination  Lacrimal gland puncture  Salivary gland puncture  Imprecise depth control | ^1–6^ |
|  | Lateral tail vein |  | < 0.1 mL | Hematoma | ^7,8^ |
|  | Lateral saphenous vein |  | 0.05-0.1 mL | Lameness  Hematoma | ^2,9,10^ |
|  | External jugular vein |  | 0.2-0.25 mL | Tissue trauma  Hemorrhage | ^11^ |
|  | “Chin puncture” aka submental route (targets vasculature running bilaterally beneath the chin, likely the inferior labial or facial veins) | ✓ | 0.1-0.2 mL | Tissue trauma | ^1,2,4,6,12^ |
|  | Retroorbital sinus aka retrobulbar route | ✓ | 0.1-0.2 mL | Tissue trauma  Hemorrhage  Ocular trauma  Corneal ulceration  Conjunctival irritation  Enophthalmos | ^3,13,14^ |
|  | Lingual vein (sublingual approach) | ✓ | 0.3 mL | Tissue trauma  Swelling of tongue  Hemorrhage  Hematoma  Aspiration of blood leading to dyspnea | ^15^ |
|  | Gingival vein (traditional) | ✓ | 0.1 mL | Bacterial contamination | ^16^ |
| **Rat** | Lateral or dorsal tail vein |  | 0.05-0.2 mL | Hematoma | ^17,18^ |
|  | Ventral tail artery |  | >1 mL | Hemorrhage  Tail tip necrosis  Thromboembolism | ^17^ |
|  | Lateral saphenous vein |  | 0.1-0.5 mL | Hematoma | ^10,19,20^ |
|  | Penile vein |  | 0.2-0.3 mL | Urine retention  Edema | ^16,21^ |
|  | Retro-orbital plexus | ✓ | 0.8-1.0 mL | Hemorrhage  Ocular injury  Orbital bone fracture | ^7,20,22^ |
|  | Lingual vein (sublingual approach) | ✓ | 1 mL | Tissue trauma  Hemorrhage  Hematoma  Swelling of tongue | ^3,23^ |
|  | Jugular vein | ✓ | 0.1-2 mL | Bruising  Hemorrhage  Infection | ^24^ |
|  | Cranial vena cava | ✓ | 0.8-2.5 mL | Hemorrhage  Cardiac injury  Vessel laceration  Tracheal injury | ^3,25^ |
|  | Gingival vein (traditional) | ✓ | 0.8 mL | Bacterial contamination | ^16^ |
|  | Gingival vein (reverse) | ✓ | 0.5-1 mL | None | Current study |
| **Guinea pig** | Lateral saphenous vein |  | 0.1-0.4 mL | Hematoma | ^10,26,27^ |
|  | Accessory cephalic vein |  | 0.1-0.4 mL | Hematoma | ^27,28^ |
|  | Ear vein |  | 0.05-0.1 mL | Hematoma | ^26^ |
|  | Jugular vein | ✓ | 1-2 mL | Hemorrhage  Tissue injury | ^26,27^ |
|  | Cranial vena cava | ✓ | 0.5-1 mL | Hemorrhage  Cardiac injury  Vessel laceration  Tracheal injury | ^28^ |
|  | Gingival vein (traditional) | ✓ | 0.5 mL | Bacterial contamination | ^28,29^ |
|  | Gingival vein (reverse) | ✓ | 0.5 mL | None | Current study |
| **Hamster** | Lateral saphenous vein |  | 0.05-0.1 mL | Hematoma | ^7,10^ |
|  | Cranial vena cava | ✓ | 0.2-1.5 mL | Tissue trauma  Vessel laceration  Tracheal injury  Well-developed clavicle may impede positioning | ^30^ |
|  | Lingual vein (sublingual approach) | ✓ | 0.75 mL | Tissue trauma  Hematoma  Hemorrhage | ^15^ |
|  | Gingival vein (traditional) | ✓ | 0.3 mL | Bacterial contamination | ^29^ |
|  | Gingival vein (reverse) | ✓ | 0.3 mL | None | Current study |
| **Chinchilla** | Cephalic vein |  | 0.15-0.5 mL | Bruising  Hematoma | ^31^ |
|  | Lateral saphenous vein |  | 0.09-0.2 mL | Bruising  Hematoma | ^31^ |
|  | Auricular vein |  | 0.15-0.5 mL | Bruising  Hematoma | ^31^ |
|  | Femoral vein |  | 0.15-0.65 mL | Hematoma  Hemorrhage | ^31^ |
|  | Ventral tail vein |  | ? | Hematoma  Edema | ^31,32^ |
|  | Cranial vena cava | ✓ | 1-2 mL | Tissue trauma  Hemorrhage  Cardiac injury  Vessel laceration  Tracheal injury |  |
|  | Transverse sinus | ✓ | ? | Bone injury  Middle ear injury | ^33^ |
|  | External jugular vein | ✓ | >1 mL | Located more medially; often cannot be visualized externally | ^27^ |
|  | Gingival vein (reverse) |  | 0.5-1 mL | None | Current study |
| **Prairie dog** | Tarsal vein | ✓ | <0.1 mL | Bruising | ^34^ |
|  | Lateral saphenous vein | ✓ | <1 mL | Bruising  Hematoma | ^34^ |
|  | Cephalic vein | ✓ | <1 mL | Bruising  Hematoma | ^34^ |
|  | Femoral vein or medial saphenous vein | ✓ | 1-3 mL | Bruising  Hematoma | ^34^ |
|  | External jugular vein | ✓ | <6 mL | Hematoma  Hemorrhage | ^34^ |
|  | Cranial vena cava | ✓ | <6 mL | Tissue trauma  Hemorrhage  Cardiac injury  Vessel laceration  Tracheal injury | ^34^ |

*Routes for which anesthesia is required to enable proper positioning and minimize the risk of complications are indicated; however, it should be noted that other routes may require anesthesia depending on patient-specific factors, such as temperament.
